# Increased signal to noise ratios within experimental field trials by regressing spatially distributed soil properties as principal components

**DOI:** 10.1101/2021.04.29.441834

**Authors:** Jeffrey C. Berry, Mingsheng Qi, Balasaheb V. Sonawane, Amy Sheflin, Asaph B. Cousins, Jessica Prenni, Daniel P. Schachtman, Peng Liu, Rebecca S. Bart

## Abstract

Environmental variability poses a major challenge to any field study. Researchers attempt to mitigate this challenge through replication. Thus, the ability to detect experimental signals is determined by the degree of replication and the amount of environmental variation, noise, within the experimental system. A major source of noise in field studies comes from the natural heterogeneity of soil properties which create micro-treatments throughout the field. To make matters worse, the variation within different soil properties is often non-randomly distributed across a field. We explore this challenge through a sorghum field trial dataset with accompanying plant, microbiome and soil property data. Diverse sorghum genotypes and two watering regimes were applied in a split-plot design. We describe a process of identifying, estimating, and controlling for the effects of spatially distributed soil properties on plant traits and microbial communities using minimal degrees of freedom. Importantly, this process provides a tool with which sources of environmental variation in field data can be identified and removed, improving our ability to resolve effects of interest and to quantify subtle phenotypes.

**IMPORTANCE:** Data from field experiments are notoriously noisy. Proper field designs with high replication aid in mitigating this challenge, yet true biological correlations are still often masked by environmental variability. This work identifies soil property composition as a spatially distributed source of variance to three types of characteristics: plant phenotype, microbiome composition, and leaf traits. We show that once identified, spatial principal component regression was able to account for these effects so that more precise estimates of experimental factors were obtained. This generalizable method is applicable to diverse field experiments.

## INTRODUCTION

Environmental variation makes the real world a noisy place to conduct science. In the context of experimental agriculture fields, variation in topography may result in uneven water moisture accumulation. Similarly, soil nutrients such as nitrogen and phosphate, are often non-uniformly distributed across a field. These unintended and often unknown sources of environmental variation may significantly affect the experimental results. The traditional approach to mitigate this variability is through experimental designs that include replicate blocks (Piepho et al. 2013; Fisher 1925). While helpful for removing variation that is relatively uniform within the blocks, true biological signal may still be masked by other experimental noise that is heterogeneous within blocks.

Analytical approaches have been used to parameterize the entire spatial variation within a field and can account for the effects on observations analytically using traditional mixed-effect modeling. These methods come in two varieties: estimating spatial-covariance structures, and spatial-smoothing using splines (Rodríguez-Álvarez et al. 2016). The former is older and canonical but more challenging to complete, and the latter is newer and easier to use courtesy of advancements in computation. Spatial-smoothing has been shown to effectively account for spatial variations in uniform barley fields and promotes genetic heritability in simulation studies (Rodríguez-Álvarez et al. 2016). While spatial-smoothing using splines does effectively address spatial variation of a trait in a field, traditional parameterizations using spatial-covariance structures do so as well and further provide intuitive metrics on the type and shape of the structure.

Spatial distribution has been considered in a variety of biological systems ranging from nematodes (Quist et al. 2019), microbiomes (Franklin and Mills 2003), forestry (Ohashi and Gyokusen 2007; Möttönen et al. 1999; Bai et al. 2012), and ionomics QTL mapping (Pauli et al. 2018). These previous studies have used spatial-covariance estimation methods to identify and associate spatial effects on various traits of interest. These methods are also the backbone of geospatial statistics where the goal is to interpolate values between sampling points (Olea 2018). These previous studies have only demonstrated the presence of spatial structure in these measurements. However, it remains a challenge to identify which factors of a multivariate dataset have an effect on the traits of interest and then adjust for the effects from all covariates while maintaining sufficient degrees of freedom for statistical inference. This challenge is similar to the challenge associated with genome-wide association studies (GWAS) that must handle population structure. In GWAS studies, phylogenetic relatedness is managed by principal component analysis (Price et al. 2006). Principal components capture axes of most variation and effectively reduce a complex multivariate dataset down to only a few independent vectors of most importance.

Here, we combine approaches from geospatial statistics and GWAS to overcome environmental variation in field studies. Specifically, we estimated spatial-covariance structures for each factor and then accounted for these effects using principal component regression. A field trial with 24 varieties of sorghum and two watering treatments, well-watered and water stressed, arranged in a split-plot design with 8 replicate blocks, was completed in 2017. Several types of data were collected including, but not limited to, plant harvest traits (height, fresh and dry weight, panicle size), soil property composition (calcium, magnesium, nitrate, organic matter, pH, phosphate, potassium, salinity, sodium, sulfate, total cations), three microbiome samplings for each plot (root, rhizosphere, and soil), leaf traits (specific leaf area, C and N content and stable isotopes of C and N), and root metabolomic profile. In this study, the soil chemical and physical properties were used as the multi-covariates that exhibited spatial-covariance structure and subsequently created micro-treatment effects throughout the field that associated with plant traits. We demonstrate that accounting for these effects via residuals of principal component regression is an effective method to improve the resolution of experimental design effects and reduce the noise caused by spatial variation within a field.

## METHODS

The field experimental design (sorghum varieties, watering regimes, and field layout) was previously described in Qi et. al., 2021. In short, we planted 24 varieties of sorghum in a split-plot design with eight replicate blocks with two watering treatments per block (well-watered and drought). End of season harvesting procedures, microbiome sampling, DNA extraction, and sequencing are also fully described in Qi et. al., 2021.

### Processing amplicon reads with VSEARCH and OTU table QC

Three microbiomes were collected for each plant: root endophytes, rhizosphere, and bulk soil, and all samples were sent for 16S PE amplicon sequencing at JGI (see Qi et.al. 2021 for extraction and sequencing methods). What follows is the VSEARCH (v2.9.0) (Rognes et al. 2016) workflow for taking the reads for each sample and processing them to curate the OTU table: merge paired ends, merge all samples, fastq filter, sequence dereplication, cluster unique sequences, remove chimeras, and read quantification. Merging paired ends had the following parameters: max diffs = 10, max diff percentage = 90, min merge length = 230, max merge length = 540. Samples were then combined into a single fasta file. Fastq filtering had the following parameters: maxee = 1, strip left = 19, strip right = 20, fastq max n’s = 0, fasta width = 0. Dereplication had the following parameters: min unique size = 1, fasta width = 0. OTU clustering had the following parameters: id percentage = 0.995, strand = both. Removing chimeras had the following parameters: fasta width = 0. Read quantification had the following parameters: id percentage = 0.9. These steps were combined all together in a directed acyclic graph (DAG) workflow and executed on a HTCondor high-throughput computation cluster. A total of 171,273 OTU’s were detected and of those 114,179 had quantification across all 1,280 samples. OTU table quality control was done in two steps: samples were removed if the total number of reads quantified across all OTU’s was less than 10,000. In addition, OTUs were removed if the total number of reads quantified across all samples was less than 100 or greater than 200,000. After applying this filter 92,385 OTU’s and 1,280 samples remained. Of the OTU’s removed only 422 had counts larger than 200,000 indicating the majority of the OTU’s removed were rare and would not have enough information to perform proper statistical analysis. Once OTU’s and samples that did not meet the quality control filters were removed, each OTU count in a sample was scaled proportionally to the same number, max number of reads per sample, so that all samples had the same number of OTU counts quantified.

### Geospatial interpolation methods

The variance of observed samples at multiple distances were calculated to produce a variogram and then, several spatial models were fitted to the data. The best fitting model was selected as the one that produced the minimal sum-of-squares errors. The spatial models that were considered include: no structure, exponential, spherical, gaussian, Matern, and Stein’s parameterization of Matern. There are three metrics that describe a spatial fit and help us understand how the sampled points are correlated. The nugget is the estimated variance between two adjacent samples and represents the noise of the data. The range is the distance at which the change in variance with respect to distance first becomes zero and represents how far away sampled points demonstrate the correlation structure. Finally, the partial sill is the variance at the range minus the nugget variance.

Each spatial structure ultimately yields estimated spatial weights as a function of the distance between two sampling points. With these weights, we can estimate values at non-sampled positions by kriging, a method to interpolate by using a weighted average of the observed values in the neighborhood of non-sampled position. We apply the ordinary kriging and let the sum of spatial weights to be one so that ordinary kriging to be unbiased (Olea 2018).

### Statistical testing for evidence of spatial structure

One major assumption of fitting spatial models is that the distribution is stationary, meaning that the mean and covariance between any two samples are the same throughout the grid. However, this field trial included two treatment factors (watering treatments and sorghum genotypes) which may have directly affected the measured soil properties. Thus, to satisfy the stationary assumption, we needed to account for any influence on the soil properties from the two treatment factors and/or their interaction. To do this, we fitted linear models including treatment factors and their interaction for each soil property separately. Residuals from these linear models correspond to soil property data after correcting for the linear effect of both treatment factors. To test for a spatial distribution, these residuals were used as the response variable of multiple models: intercept-only, and others with different spatial covariance structures (spherical, exponential, gaussian, linear, and rational-quadratic). The likelihood of the intercept-only model was compared to the likelihood of each spatial model using a likelihood ratio test. A soil property is considered to exhibit evidence of spatial structure if any of five spatial covariance structure models are significantly more likely than the intercept-only model

### Soil property composition sampling and processing

Selected plants were excavated using a shovel to a depth of 12 – 14 inches. The soil (approximately 200 g) from the excavated root ball was shaken off into a wash pan in the field, homogenized and collected into a quart-size Ziploc bag. In addition to the collection of roots for microbiome analysis a subset of roots were collected for metabolite analysis as described in (Sheflin et al. 2019). The soil used for chemical and physical analysis was stored in the Ziploc bags at 4°C and sent to Ward labs for analysis of pH, buffer pH, sum of cations (CEC), base saturation (%), soluble salts, organic matter, nitrate-nitrogen, phosphorus, potassium, calcium, magnesium, sodium, sulfur, zinc, iron, manganese and copper.

### Root metabolomics sampling and processing

#### Non-targeted metabolite profiling using gas chromatography mass spectrometry (GC-MS)

Metabolite extraction was conducted by weighing out 19 - 21 mg of each freeze-dried sorghum roots and placing them into clean 2 mL autosampler glass vials (VWR, Radnor, PA, USA). Automated control of sample extraction (i.e., solvent proportions, solvent volumes, sample agitation and supernatant transfers) was accomplished using a standalone Gerstel MultiPurpose Sampler (MPS). Samples were extracted by adding 770 μl of methyl-tert-butyl-ether (MTBE) and 385 μl to each vial and vortexing on the MPS at room temperature for 30 min. To separate organic and aqueous layers, 640 μl of water was added to the remaining extract and vortexed for 15 min. Samples were then centrifuged for 25 min at 3500 rpm at 4 C. The organic layer was extracted twice by transferring into a new 2 mL autosampler vial without disturbing the lower layer then adding 600 μl of MTBE and transferring again. The aqueous layer was also extracted twice by transferring out of the vial into a new 2 mL autosampler vial without disturbing the pellet then adding 300 μl of methanol and 300 μl of acetonitrile, vortexing for 3 min and transferring again. The aqueous layer was completely dried under N gas, resuspended in 300 μl of 75% methanol. 20 μL of the aqueous layer from each sample was transferred to another set of glass vials, centrifuged for 2 min at 3500 rpm and then dried under N_2_ (g) for 30 min. Dried samples were stored at −80 °C until derivatization. Derivatization (methoximation and silylation) took place immediately prior to running the samples. Dried down samples were allowed to warm to room temperature and then re-suspended in 50 μL of methoxyamine HCl (prewarmed to 60 °C) and centrifuged for 30 sec. Samples were then incubated at 60 °C for 45 min, followed by a brief vortex, sonication for 10 min and an additional incubation at 60 °C for 45 min. Following this, the samples were centrifuged before receiving 50 μL of N-Methyl-M (trimethylsilyl) trifluoroacetamide (MSTFA) + 1 % trimethylchlorosilane (TMCS) (ThermoFisher Scientific, Waltham, MA, USA), briefly vortexed and incubated at 60 °C for 40 min, as described previously (Chaparro et al. 2018). Samples were loaded (~100 μL) into glass inserts within glass autosampler vials and centrifuged for 30 sec prior to GC-MS analysis. In addition, a pooled extract was created by combining equal volumes of each sample into one glass vial for use as a consistent representative quality control sample (QC).

GC-MS analysis was performed using a Trace 1310 GC coupled to a Thermo ISQ mass spectrometer (ThermoScientific). Derivatized samples (1 μL) were injected in a 1:10 split ratio. Metabolites were separated with a 30-m TG-5MS column (Thermo Scientific, 0.25 mm i.d. 0.25 μm film thickness). The GC program began at 80 °C for 0.5 min and ramped to 330 °C at a rate of 15 °C per minute and ended with an 8 min hold at a 1.2 mL · min^−1^ helium gas flow rate. The inlet temperature was held at 285 °C and the transfer line was held at 260 °C. Masses between 50-650 m/z were scanned at five scans/sec after electron impact ionization.

Metabolomic data processing was conducted as previously described (Chaparro et al. 2018). GC-MS files were converted to .cdf format and processed by XCMS in R (Smith et al. 2006; Mahieu et al. 2016; R Core Team 2015). All samples were normalized to the total ion current (TIC). RAMClustR was used to deconvolute peaks into spectral clusters for metabolite annotation (Broeckling et al. 2014). RAMSearch (Broeckling et al. 2016) was used to match metabolites using retention time, retention index and matching mass spectra data with external databases including Golm Metabolome Database (Hummel et al. 2007; Hummel et al. 2013) and NIST (Broeckling et al. 2016).

### Leaf traits analyses

The middle portion (10-12 cm long) of the uppermost fully expanded leaf from individual plants was harvested in a coin envelope for the analysis of specific leaf area, C and N content, and stable isotopes of C and N. Leaf samples were oven-dried at 65 °C to a constant mass and ca. 2.5 mg of the dry leaf was subsample using a custom-made leaf punch system in a tin capsule. The leaf punch provided the leaf area of subsample, which were weighed to estimate specific leaf area (SLA). The N, C and δ^15^N and δ^13^C concentrations of dry leaf were determined by combusting encapsulated samples in an elemental analyzer (ECS 4010, Costech Analytical Technologies) coupled to a continuous flow isotope ratio mass spectrometer (Delta XP, Finnigan MAT) at the Stable Isotope Core Laboratory, Washington State University.

## RESULTS

### Geospatial statistics interpolates soil property composition throughout the field

We conducted a field experiment in which sorghum and its associated microbiome were evaluated across two different watering treatments (Qi et. al. 2021). As is typical of field studies, the collected data showed significant variability across all measured parameters (biomass, leaf traits, metabolites, and microbiome (root, rhizosphere and soil)). In this study, we also measured several different soil properties at multiple points throughout the field including: organic matter, pH, phosphate, nitrate, sum of cations, calcium, magnesium, potassium, sodium, sulfate and salinity compositions. We hypothesized that variation across the field in these soil properties may explain some of the variability in the other measurements.

First, because only a limited set of points across the field were sampled for soil properties, there was a need to estimate the missing values (Figure 1A). We hypothesized that sample proximity would correlate with the measured soil properties. To assess this type of spatial correlation, we employed techniques from geospatial statistics to capture the correlation structure of any pair of samples in the field. Of the twelve properties tested, six properties exhibited evidence of spatial distributions (see methods) (p-value < 0.05): salinity (mmho), nitrate (ppm), sulfate (ppm), calcium (ppm), magnesium (ppm), phosphate (ppm). For these six soil properties we estimated the missing values throughout the field. Interpolation of values between sampling points was performed by leveraging spatial correlation structure to predict unobserved values, a process called kriging (see methods) (Supp Table 1). To test the kriging accuracy, we performed a leave-one-out cross validation for each soil property. Through this analysis we observed that the error of the predictions, when scaled to unit variance of the observations, exhibit distributions that resemble the expected standard Z-distribution (Supp 1A). The ratio of the partial sill to the nugget, see methods for definitions, is a proxy for the magnitude of the variance that is attributable to the spatial structure. Comparing this ratio for nitrate and phosphate, shows that the phosphate spatial correlation was much stronger than the nitrate spatial correlation (Fig 1B,C). Calcium also exhibited correlation structure of distances larger than nitrate, but much smaller than phosphate (Fig 1D). To visualize the spatial structure for each soil property, the kriged values of each property were centered around the mean and scaled to unit variance (Fig 1A-D). This analysis revealed that the soil properties exhibited different topographies across the field. For example, phosphate levels were high in a band across the center of the field while nitrate and calcium levels were more variable with several high and low spots (Fig 1B-D). We also considered correlation between the different soil properties and observed several correlation blocks, implying similar spatial structures (Supp 2A).

**Figure 1.**
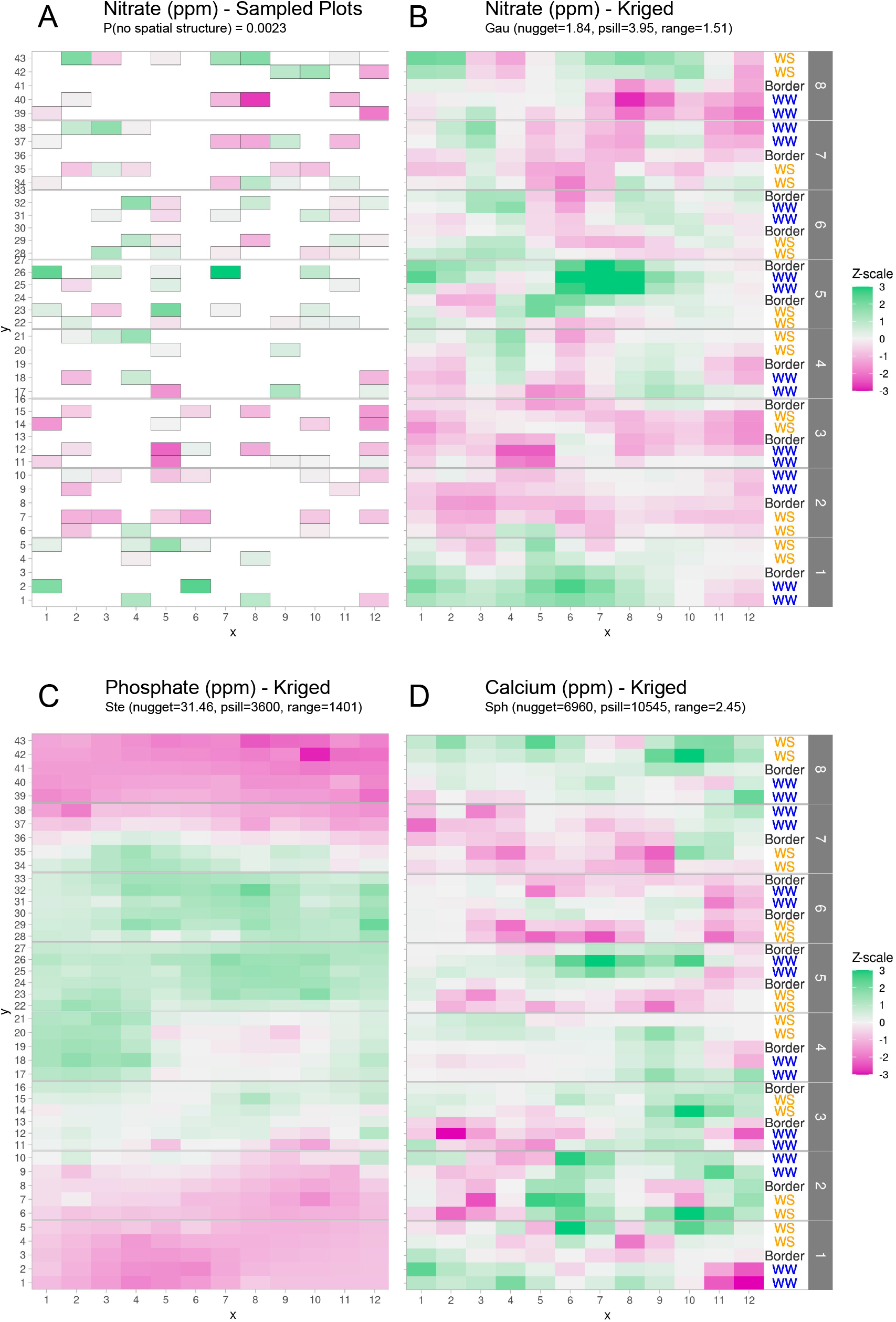
Graphical depictions of field layout where each cell is a plot in the field. Water treatment is specified on right; WS = water stressed, WW = well watered. Eight split-plot replicate blocks are denoted in gray vertical bars. Color scale represents data with genotype and treatment removed. Green indicates larger than average, white indicates approximately average, and magenta indicates below average values. (A) Nitrate values are shown for each cell (outlined in grey) that were sampled for soil property analysis. (B) kriged nitrate values to estimate nitrate levels in unsampled plots (C, D) Kriged values for phosphate and calcium. Variogram fit of spatial model is indicated with model type, nugget, partial sill, and range.

### Soil property variation influences plant phenotypes and microbiome composition

The above analyses clearly showed that soil properties were variable across the field site. However, it was not clear whether the observed variation was large enough to affect plant associated phenotypes or the microbiome. To address this, we used constrained analysis of principal coordinates (CAP). With CAP, it is possible to identify specific effects on a multidimensional dataset while acknowledging variation due to other effects. For example, to understand if and how soil property variation affected microbiome composition, we first had to control for the effects of the watering treatments, the different genotypes and their interactions. A permutation ANOVA, using 999 iterations, was performed to assign statistical significance to the kriged properties. CAP and PERMANOVA were done for each of the three microbiome compartments (root, rhizosphere, and soil), and association was assessed for each soil property measured. From this analysis we observed that the root microbiome was invariant to all soil property variations. In contrast, the rhizosphere and soil microbiomes were influenced by the variation in several soil properties (Fig 2A). Additionally, there were some soil properties (salinity, sulfate, and calcium) whose variation affected either the rhizosphere or soil microbiomes but not both. This suggests that microbiome compartments are differentially sensitive to different types of soil property variation. CAP was also applied to annotated root metabolomic profiles. In contrast to the large effects seen in the microbiomes, no soil property variances were associated with changes in the metabolomic profile (Supp 3A). This may reflect the relative stability of the metabolites identified from GCMS which mostly represents primary metabolites, or the sensitivity of measurements.

**Figure 2.**
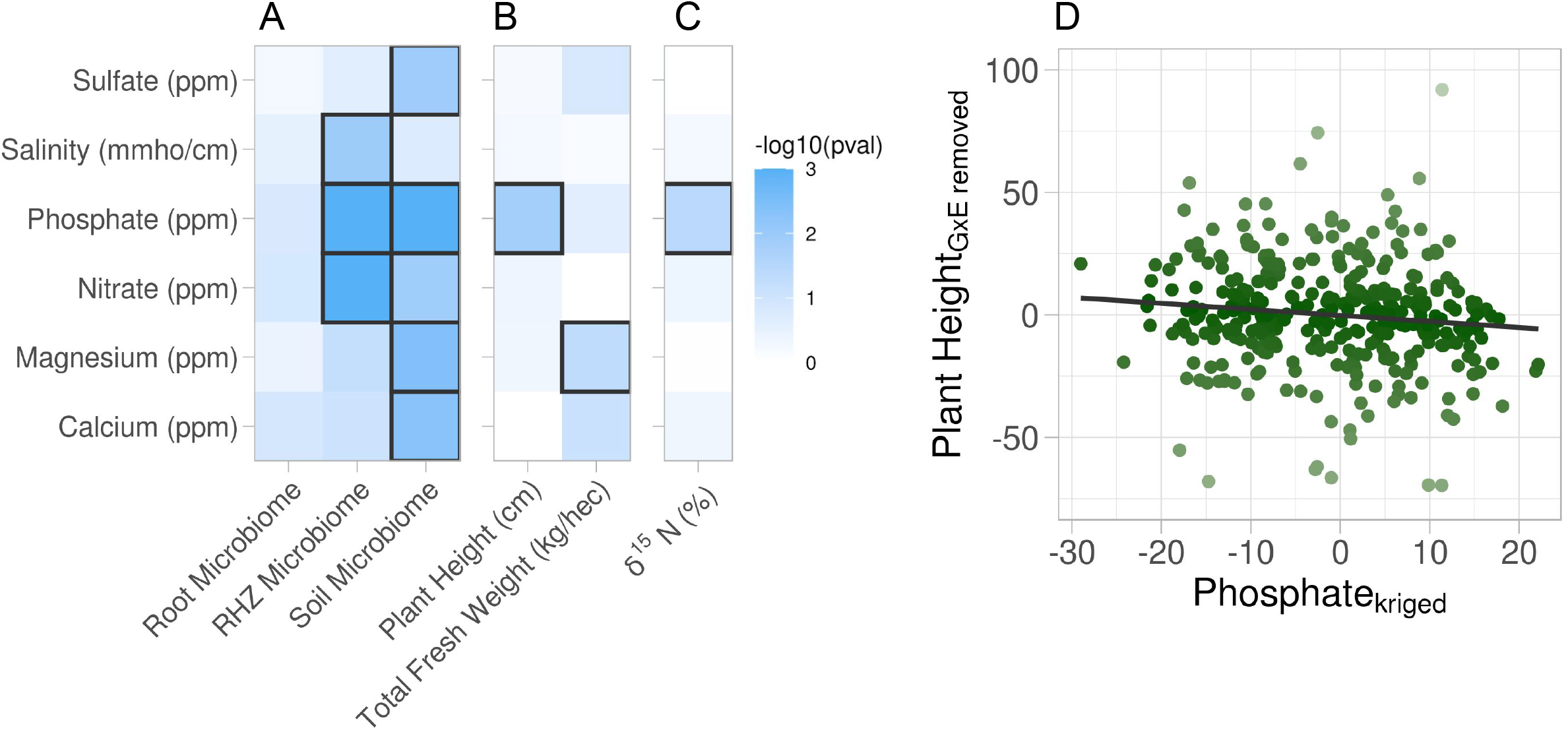
Association of soil property variations with multiple phenotypes. Six soil properties were assessed for effect on root microbiome and plant phenotypes using permutation ANOVA. Cells are colored by −log10 p-value of the effect. (A) Effect of each soil property on microbiome beta diversity from three root compartments: root (endosphere), RHZ (rhizosphere) and soil (bulk soil), while constraining on genotype and treatment. (B,C) Effect of each soil nutrient on the height and weight (B) and leaf δ^15^N (C) using type III sum of squares while including treatment, genotype, and interaction as additional fixed effects. (D) Example effect of kriged phosphate, axis, on plant height, adjusted for genotype and treatment, y-axis.

While microbiome and metabolite data are highly multivariate, plant phenotypes and leaf traits are much less so and are therefore suited to univariate statistical analyses. Just as the microbiome has the potential to be influenced by the soil property variations, the same could be true for the univariate phenotypes. Similarly, the effects due to the experimental design must be acknowledged to more precisely evaluate the association a given property has on the phenotype. Mixed effect models with random effect for the split-plot replicates and multivariate-normal spatial correlation structure were created for each soil property-phenotype pair. The precise effect a given soil property has on a phenotype was evaluated using type-III sum of squares to account for the other sources of variance (treatment, genotype, and the interaction) on the phenotypes. Traditional F-statistics from the analysis of variance (ANOVA) tables revealed that plant height and total fresh weight are influenced by soil phosphate and magnesium variation, respectively (Fig 2B). Similar modeling of the leaf traits indicated the soil phosphate variation is significantly associated with δ^15^N (Fig 2C). Many other phenotypes were examined and did not have statistically significant associations with the variation in soil properties (Supp 3A). Closer examination showed that soil phosphate levels are mildly inversely correlated to plant height (Fig 2D). This supports the hypothesis that excess phosphate inhibits plant growth and development (Shukla et al. 2017; Song et al. 2016) and suggests that the levels found in the center of the field were too high for optimum sorghum growth.

### Statistical approach to place field noise into principal components

We have shown that many of the soil properties exhibit spatial distribution and influence various plant and microbiome traits. Therefore, to understand the effects of treatment and genotype on these traits, the effects of the soil properties must first be accounted for. The replication in our study was not sufficient to include all the soil properties as covariates to account for their influence -- this would require a degree of freedom for every soil property. A generalized approach to overcome limited sample size is reducing the dimensionality of the covariates using principal component analysis and regressing against the first several principal components, known as principal component regression. In this approach, the principal components retain a percentage of the influence from the individual properties and can be used as a proxy to adjust for as much variation as possible.

As above, our goal was to create spatial models using the generated principal components and so the stationary requirement must be met. Using only the observed data, not the kriged values, we first adjust for treatment, genotype and interaction effects from each soil property by fitting linear models and obtaining model residuals, and the residuals were then processed with principal component analysis. To test the success of accounting for our experimental design, we investigated clustering within the first two PCs and observed that indeed the treatment and genotype effects were accounted for (Fig 3A). We selected the first three components as regressors which represented approximately 66% of the total variance in the soil property data (Fig 3B). Next, we visualized the contribution of each soil property within each principal component (PC) (Fig 3C). PC1 is primarily represented by calcium, magnesium, potassium, sodium and sum of cations. PC2 is primarily represented by sulfate, salinity and pH. PC3 is primarily represented by phosphate and nitrate. Then, we used kriging, as described above, to interpolate the missing values for the rest of the field. Since many soil properties exhibited spatial distributions (Fig 1), we expected that the principal components would also display a spatial distribution. Indeed, the spatial distribution of the kriged first principal component resembles the calcium distribution which emphasizes the contribution of that property (Fig 1D and Fig 3D). In summary, through this approach, we revealed specific sources of field level noise and successfully captured a significant proportion of the field level variation into three principal components.

**Figure 3.**
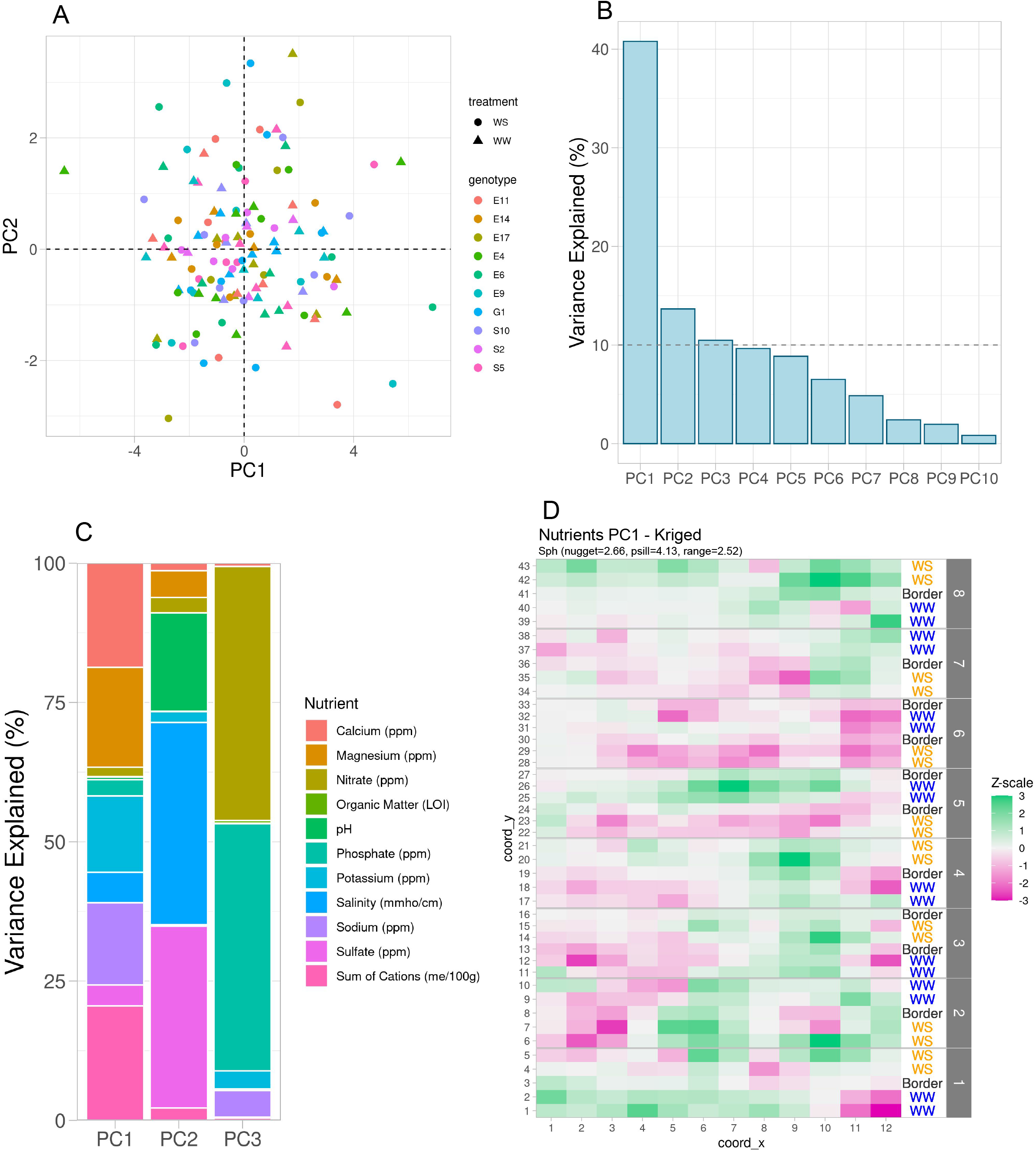
Variation in soil properties can be collapsed into principal components. (A) First two principal components of soil property residuals as x-axis and y-axis respectively colored by genotype and shaped by treatment. (B) Skree plot of the first 10 principal components. Shown is the percent variance explained of the total property variance by each component. Dashed line is at 10% variance explained. (C) For the first three components, colored is the contribution of each soil property to its respective variance explained within each component. (D) Spatial distribution of kriged PC1. Each cell colored by scaling the values to unit variance. Variogram fit with nugget, partial sill and range displayed.

### Using principal components to de-noise field data

As a final step, we sought to address the variation associated with the three principal components from the plant and microbiome data. Starting with the microbiome data, generalized mixed-effects models based on zero-inflated negative-binomial distribution were created for each individual microbe where the microbe counts were the dependent variable, the first three principal components of the soil property variations were fixed effects, and random intercepts for each split-plot replicate each having multivariate normal spatial correlation structure. An adjusted count for each microbe was created by dividing the raw count by the corresponding estimated scale effect of the three principal components, and the intercept according to the fitted generalized linear model. Since we accounted for a source of variation that is invariant to the microbiome compartments, we expected the variance within each of the compartments to decrease and the difference between compartments to be more obvious. To test this, we combined the original observed counts and the adjusted counts for these remaining microbes and performed principal component analysis. This revealed clustering that demonstrated larger distances between microbiome compartments using the adjusted counts versus the observed counts. This indicates the sources of variation from the soil properties were better controlled thereby increasing the differentiation between compartments. Within each compartment, the root microbiome was least affected by soil factors which was demonstrated by the small distance between the observed and adjusted counts, followed by rhizosphere with a larger separation, and soil being the farthest and most affected (Fig 4A). The adjusted counts produce clusters that are larger than their respective original counts again indicating sources of variation were addressed so that the within compartment effects are better elucidated (Fig 4A). Next, we sought to identify treatment effect changes within each of the tissue compartments. Given the strong compartment differentiation, identifying clusters within a compartment required principal component analysis to be performed on each compartment individually. In the rhizosphere we observed treatment differentiation using both the observed and adjusted counts; however, the distance between the centers of each cluster is larger after adjustment further indicating within-group variation being reduced (Fig 4B).

**Figure 4.**
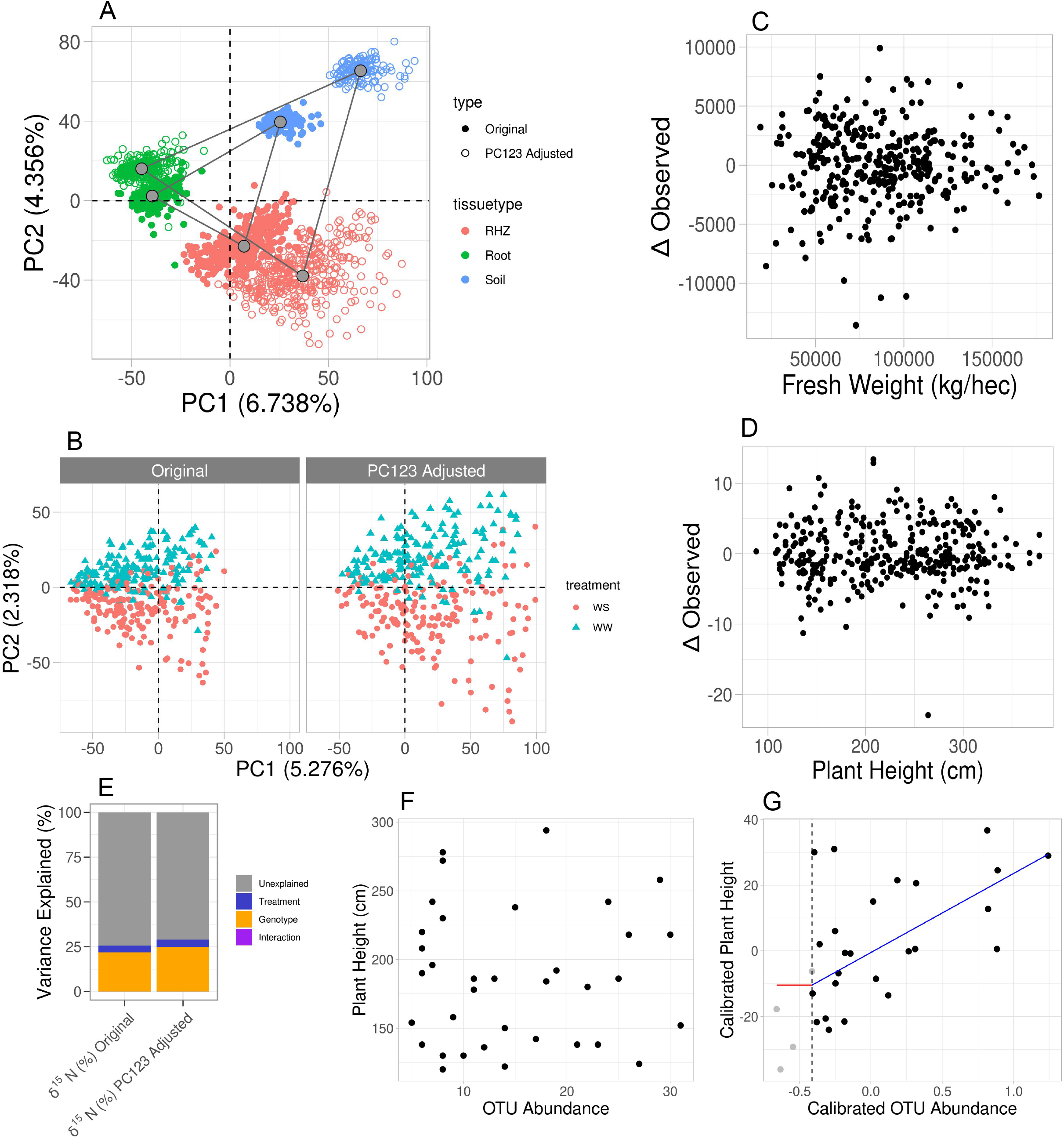
Accounting for influence from soil property variance within microbiome data reveals plant phenotypes that correlate with microbe abundance. (A) Principal component analysis on the combined raw and residual microbiome tables. Shown are the first two components with their respective variance explained. Samples are colored by tissue type and shaped by original or residual values. Gray points are the centers of each respective cluster, and gray lines connect the centers of each cluster. (B) Principal component analysis on the rhizosphere samples only using the combined raw and residual microbiome tables. Left panel uses the original counts and right uses adjusted counts as described (methods). Each panel has colors corresponding to the treatment for each sample. (C) Observed plant height values, x-axis, and the change in that value as a result of the adjustment, y-axis. (D) Similar to C, shown are the fresh weight values and their respective changes. (E) Partial correlations of experimental design variables in leaf δ^15^N before and after principal component regression. (F,G) For only water-stressed samples and only the soil microbiome, plant height, y-axis, and OTU abundance of *Microvirga*, x-axis, before and after principal component regression. Shown in G, is the fit of a change-point model where the red line is no change before threshold, the vertical dashed line, and the blue line is a linear fit after the threshold.

To assess how principal component regression affected the plant phenotypes, we compared the data before and after signal marginalization. Prior to removing noise from soil properties, plant height and fresh weight both showed drought effects, and after performing principal component regression, these effects were maintained (Supp 4). Further, by plotting the original values against the change for each value after adjustment, we showed that our correction method is equitable for all plant sizes; in other words, short and tall plants were not overly adjusted either positively or negatively (Fig 4 C, D). Additionally, the variation attributable to soil properties was as much as +/− 10% for fresh weight and +/− 6% for plant height (determined by the ratio of the standard deviation of the change to the standard deviation of the observations). This indicates that if an experimental factor is expected to have an effect size less than the soil property effect size, the contribution of the soil property variations may mask the ability to resolve the effects of the experimental factor. For instance, leaf delta δ^15^N showed an association with phosphate and calculated partial correlations for the treatment, genotype and interaction effects on this trait and found that the difference in R^2^ between the adjusted and unadjusted is approximately 0.05 (Fig 4E). This shows that the soil properties explain about 5% of the total variation in δ^15^N, which is similar to variance explained in both plant height and fresh weight.

Microbiome analyses often have less-than-ideal replication and therefore elimination of confounding variations is crucial in identifying effects of interest, such as identifying plant-growth promoting microbes. For instance, change-point models can identify microbial impacts on plant phenotypes once the abundance of the microbe reaches a particular level (Qi et. al. 2021). We show that in water stressed samples before the adjustment on both plant height and microbe abundance of *Microvirga* in the soil microbiome, we do not see any evidence of an association but after accounting for the variance from the soil properties there is a strong plant-growth association (Fig 4F, G). This further emphasizes that if an experimental factor is expected to have an effect size that is less than the soil property effect size, the experimental factor would be lost to noise. This result demonstrates that even though these are relatively small changes to each of these data types, when combined for associations, the effect can be large.

## DISCUSSION

Large scale trials within complex environments are an important component of many biological subdisciplines. Because of environmental variability, these experiments must include high levels of replication and even so, results often fail to repeat in subsequent trials. For agricultural field studies a major source of variation is heterogeneous soil property distributions that create their own micro-treatments and are covariates to planned experimental designs. Because these micro-treatments are often unknown, and therefore not accounted for, they show up as experimental noise and may lead to false positives. For instance, nitrogen is known to affect plant growth (size, color, yield, etc) (Chapin et al. 1987, Veley et al. 2017). If nitrogen is unevenly distributed across a field experiment aimed at characterizing biomass among diverse genotypes, the variability in nitrogen may confound the experiment. In this study, by intentionally measuring multiple soil properties across the field experiment, we were able to account for this known variation through principal components and gain novel biological insight into field relevant interactions between plants and microbes (Qi et al. 2021).

While the approach described here represents a major advance forward, we acknowledge several opportunities for further improvement. For example, the soil property data was collected approximately one month prior to harvest and could have been impacted by a multitude of things in that time. Future studies might gather soil property composition at multiple time points, including before planting, to generate paired data, and of course at the time of phenotyping. We predict this approach would still be able to account for soil property variation under the assumption that the soil properties themselves are relatively stable. Additionally, advancements have been made in spatio-temporal modeling using Bayesian hierarchical modeling with time as an autoregressor (Finley et al. 2015; Rushworth et al. 2014) which may prove powerful if soil property composition were densely sampled over time. We also note that replication remains a crucial aspect of these types of experiments. In this analysis, we lost three degrees of freedom by using the first three principal components (PC) of the soil property data in a regression to account for their contributions on the phenotypes. Had a fourth PC shown a significant source of variance, degrees of freedom would have become limiting. On the other hand, had we included additional replication, it may have been possible to correct for covariates such as the soil properties by directly regressing on the properties themselves, rather than using a dimension reduction technique.

We note that in these datasets, the stable isotopes and primary metabolite profiles are invariant to the measured soil properties. For the isotopes, this may be indicative of stability relative the fluctuations in the properties across the field but it is possible that if the soil property variations were larger, then a relationship might be established. The metabolite profile used in these analyses are only those captured from GC/MS and mostly consist of primary metabolites such as sugars, organic and amino acids, small phenolics, and fatty acids. It may be true that secondary metabolites that were not examined in these analyses may associate with the soil property variations. We observe that a relatively small amount of variation in plant height and weight are attributed to the soil property composition, other types of data, particularly microbiome composition, are much more susceptible. Microbiome quantification has been shifting from using operational taxonomic units (OTUs) to amplicon sequence variants (ASVs) which are designed to identify and retain more specific bacterial identification. Some microbes were overtly sparse across the samples and principal component regression could not successfully estimate model parameters. The microbiome table generation pipeline used for this field allows for the identification of very sparse microbes by way of using 99.5% identity clustering which results in 23,617 OTUs detected. After applying principal component regression, approximately 25% of the microbes were successfully modeled and retained. The methods proposed here should also be applicable to those types of tables as well with one caveat: ASV tables are sparser than OTU tables. While the methods proposed here are zero-inflated, likely the percent of ASVs that would not be successfully modeled would be larger than what would be observed with an accompanying OTU table of the same data.

In conclusion, here we demonstrated the impact of spatially distributed soil property variations on several phenotypes of interest and present principal component regression as a method to alleviate the effects analytically. Phenotypes range in their sensitivities to the soil properties and could contribute large amounts of variation to the recorded observations leading to false-positive or false-negative results. In this field study, the microbiome communities were identified to be heavily influenced by the soil properties while plant phenotypes were more resilient but nonetheless affected. Identifying sources of variation and removing their influence enhances the ability to resolve other effects of interest and enables more honest, reliable, and believable quantification of subtle phenotypes.

## SOFTWARE USED AND DATA AVAILABILITY

All analyses herein were performed in R using the following packages: raster(3.4.5), ggplot2(3.3.3), deldir(0.1.21), vegan(2.5.5), plyr(1.8.4), gridExtra(2.3), reshape(1.4.3), FactoMineR(2.4), factoextra(1.0.7), chngpt(2019.3.12), stringr(1.4.0), gstat(2.0.6), sp(1.4.2), scales(1.0.0), lme4(1.1.21), nlme(3.1.140), parallel(3.5.2), patchwork(1.1.1). All data and scripts used to create all figures and perform all analyses can be found at 10.5281/zenodo.4715924.

---

**Supplemental 1.**
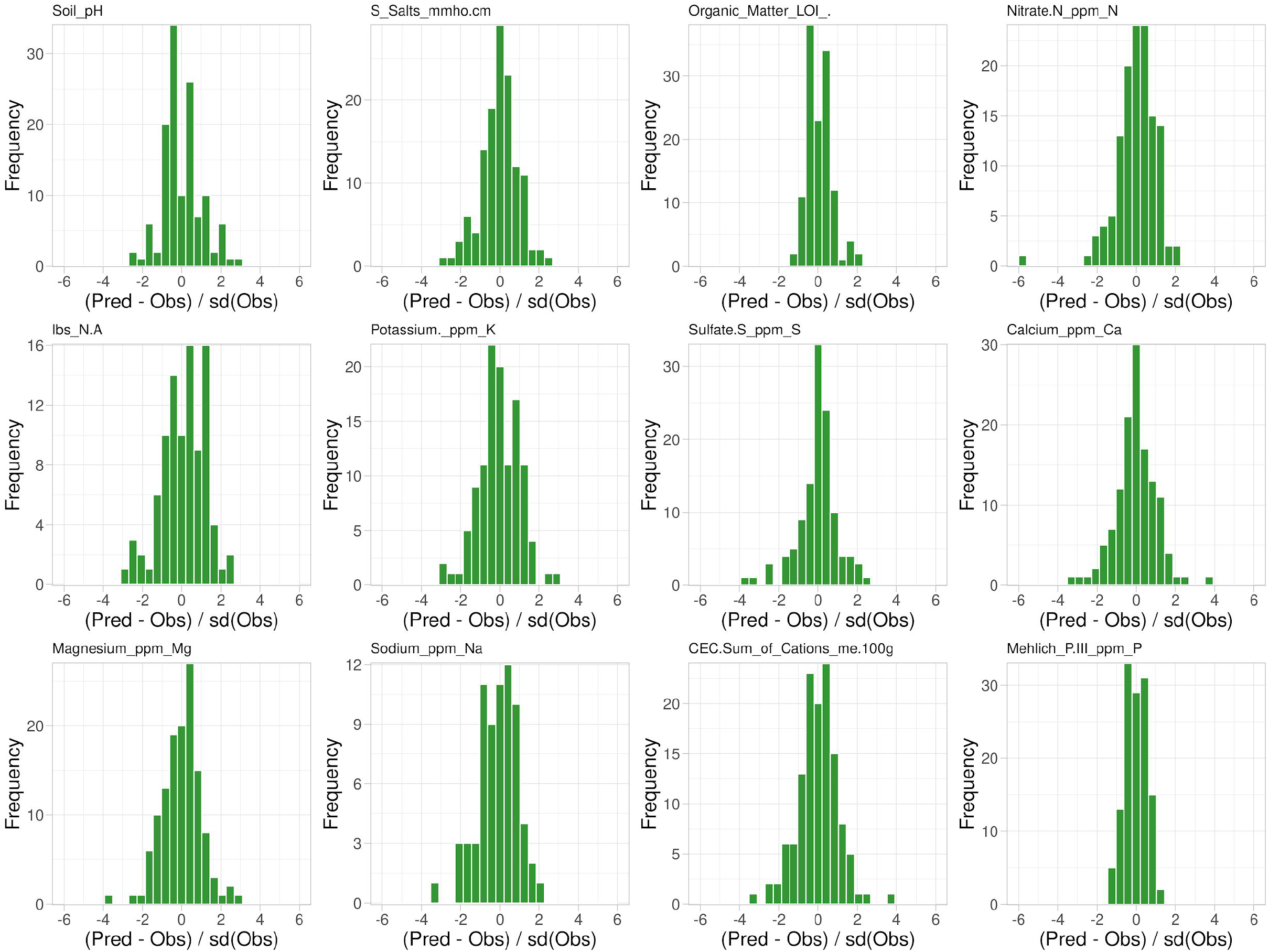
(A) Leave-one-out cross-validation of soil property observations. Shown is the difference between the predicted and the observed values for each sample normalized to the standard deviation of the observations, x-axis, and their respective frequency, y-axis.

**Supplemental Table 1.** Shown are sum-of-square errors for each soil property (columns) and each spatial model tested (rows). Cells that are colored green are those models that have minimal errors and are chosen to be the best fit model for kriging for the respective soil property.

|  | Salinity | Nitrate | Sulfate | Calcium | Magnesium | Phosphate |
| --- | --- | --- | --- | --- | --- | --- |
| Nugget Only | 1.92115E-05 | 203.668 | 10645.6 | 833211000 | 2261360 | 149688 |
| Exponential | 1.1769E-06 | 41.2651 | 1909.53 | 17886300000 | 65136300 | 27556.2 |
| Spherical | 2.03099E-06 | 25.5247 | 2058.42 | 244155000 | 1038010 | 32351.9 |
| Gaussian | 2.16269E-06 | 22.3183 | 289060 | 17669600000 | 64365000 | 100531 |
| Matern | 1.13273E-06 | 23.0069 | 1682.03 | 266373000 | 1087980 | 18961.6 |
| Stein's Matern | 1.13273E-06 | 23.0069 | 290357 | 17886300000 | 65136300 | 18827.8 |

**Supplemental 2A.**
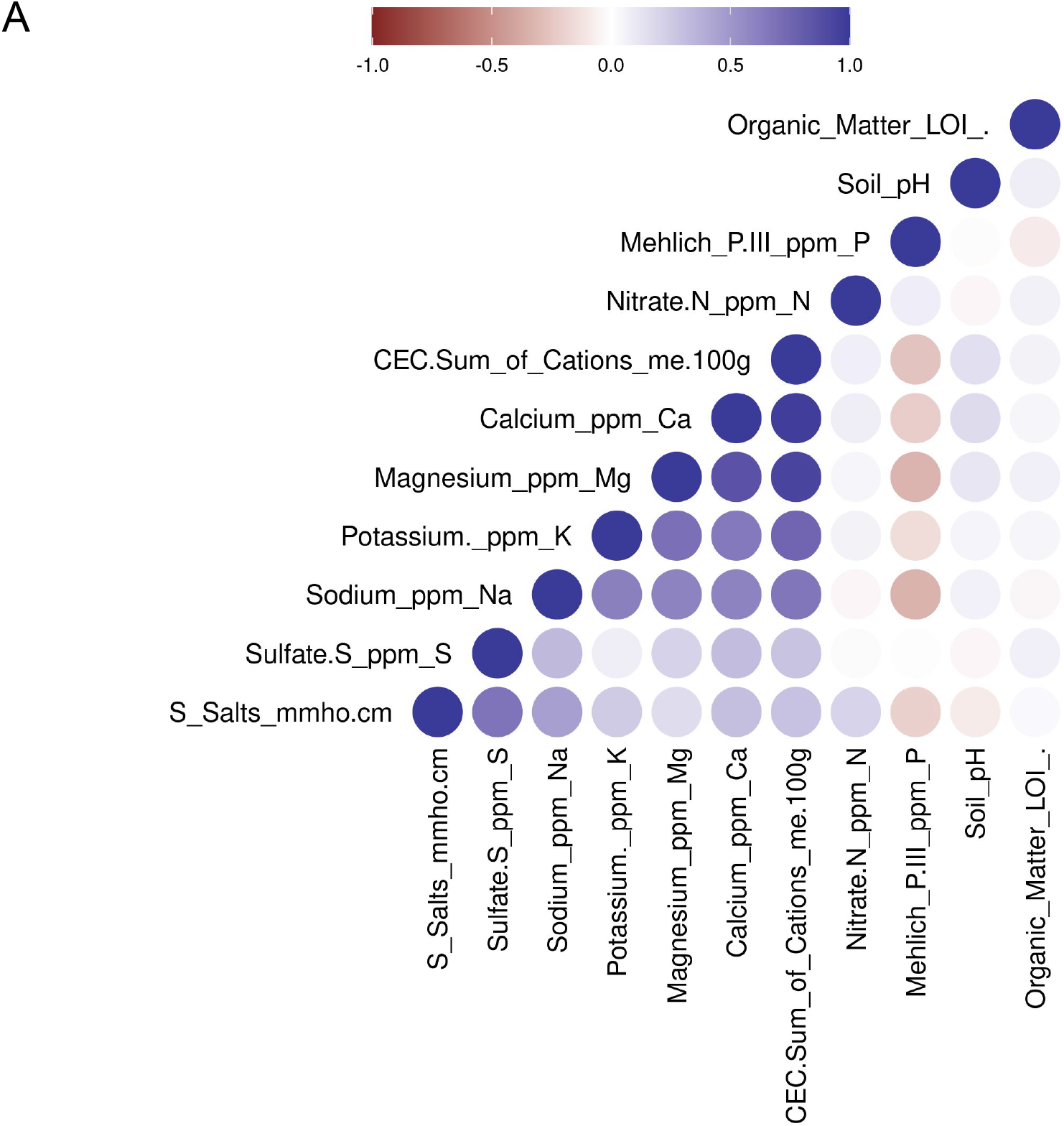
(A) Pearson correlations between all pairwise soil properties. Negative correlations are depicted as shades of red, and positive correlations are depicted as blue. Soil properties are ordered by hierarchical clustering based on euclidean distance and complete agglomeration.

**Supplemental 3A.**
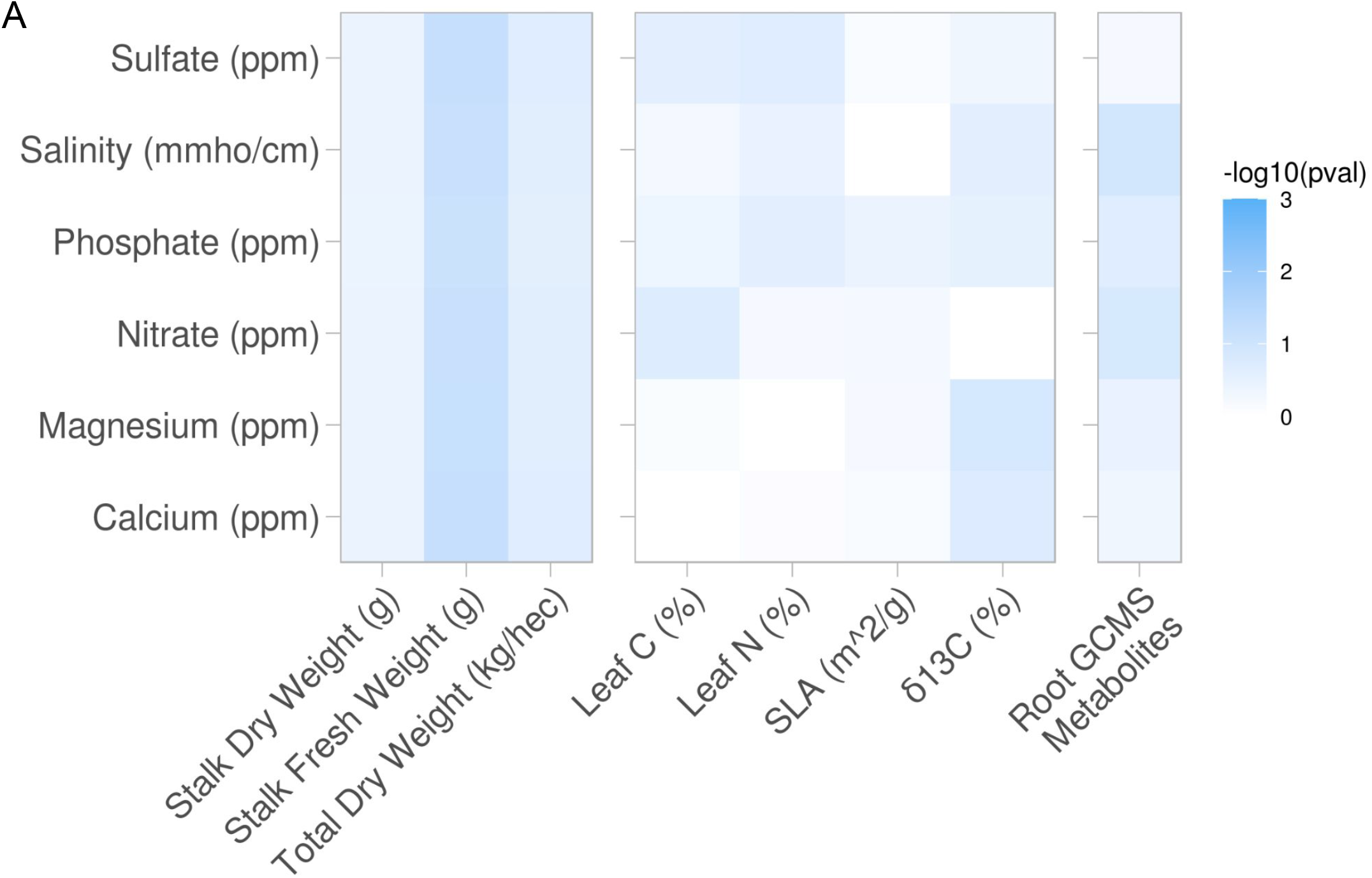
Same analysis as Fig 2 A,B,&C, but here are shown the phenotypes that did not demonstrate soil property associations.

**Supplemental 4.**
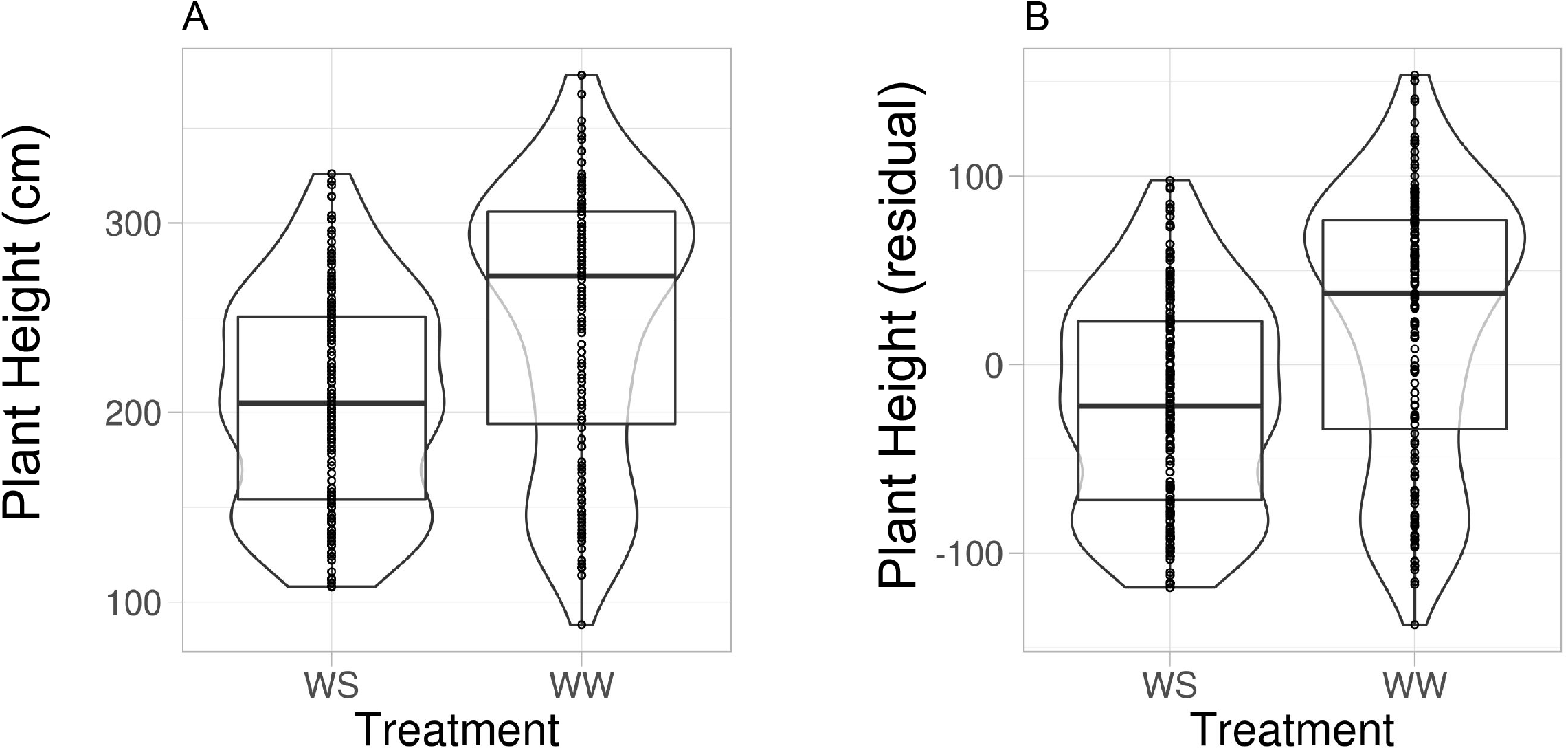
Effect of applying principal component regression on plant morphology. (A) Boxplots, overlaying violin plots, of plant height, y-axis, against the watering treatments, x-axis (B) Same as A, but displaying the residuals of plant height, y-axis, from principal component regression model using dimension reduced soil properties as covariates. See (Qi et al. 2021) for additional details and analysis on these data.

